# Temporal and sex-dependent gene expression patterns in a renal ischemia-reperfusion injury and recovery pig model

**DOI:** 10.1101/2021.09.01.458480

**Authors:** Stéphane Nemours, Luis Castro, Didac Ribatallada-Soriano, Maria Eugenia Semidey, Miguel Aranda, Marina Ferrer, Alex Sanchez, Joan Morote, Gerard Cantero-Recasens, Anna Meseguer

**Affiliations:** Renal Physiopathology Group, CIBBIM-Nanomedicine, Vall d’Hebron Research Institute, Barcelona, Spain; Biomedical Research in Urology Group, Vall d’Hebron Research Institute, Barcelona, Spain; Department of Pathology, Hospital Vall d’Hebron, Barcelona, Spain; Rodent Platform, Vall d’Hebron Research Institute, Universitat Autònoma de Barcelona, Barcelona, Spain; Unitat d’Estadística i Bioinformàtica, (UEB), Vall d’Hebron Research Institute, Barcelona, Spain; Departamento de Genética, Microbiología y Estadística. Universitat de Barcelona, Barcelona, Spain; Departament de Bioquímica i Biologia Molecular, Unitat de Bioquímica de Medicina, Universitat Autònoma de Barcelona, Bellaterra; Spain; Red de Investigación Renal (REDINREN), Instituto Carlos III-FEDER, Madrid, Spain

**Keywords:** kidney, ischemia-reperfusion, pathway enrichment, sex difference, pig model, gene expression patterns

## Abstract

Men are more prone to acute kidney injury (AKI) and chronic kidney disease (CKD), progressing to end-stage renal disease (ESRD) than women. Severity and capacity to regenerate after AKI are important determinants of CKD progression, and of patient morbidity and mortality in the hospital setting. To determine sex differences during injury and recovery we have generated a female and male renal ischemia/reperfusion injury (IRI) pig model, which represents a major cause of AKI. Although no differences were found in blood urea nitrogen (BUN) and serum creatinine (SCr) levels between both sexes, females exhibited higher mononuclear infiltrates at basal and recovery, while males showed more tubular damage at injury. Global transcriptomic analyses of kidney biopsies from our IRI pig model revealed a sexual dimorphism in the temporal regulation of genes and pathways relevant for kidney injury and repair, which was also detected in human samples. Enrichment analysis of gene sets revealed five temporal and four sexual patterns governing renal IRI and recovery. Overall, this study constitutes an extensive characterization of the time and sex differences occurring during renal IRI and recovery at gene expression level and offers a template of translational value for further study of sexual dimorphism in kidney diseases.

**AUTHOR SUMMARY:** **Kidneys’ correct functioning** is essential for optimal body homeostasis, being their basic functions blood filtration and excretion of wastes and toxins. Inherited or acquired conditions can cause renal dysfunction requiring renal replacement therapy, which will affect patients’ life quality and survival. A major cause of kidney failure is **the renal ischemia/reperfusion injury** (IRI), which occurs in many clinical situations like kidney transplantation or aortic aneurysm surgery. Interestingly, men are more susceptible to IRI than women, being women more protected against kidney injury. However, the genetics regulating these sex differences in injury and renal repair remained unknown.

Here, we provide a novel porcine model to study renal injury and recovery in both males and females. Using this model, we have identified the gene sets involved in renal injury and recovery processes. Moreover, global genetic analyses allowed us to discover the temporal and sex-dependent patterns that regulate those gene sets and, finally, kidney damage and repair. A relevant finding of our study is that males develop a feminized genetic profile during recovery, which may represent a survival mechanism to diminish the androgenic pro-damage effects on kidney cells. To sum up, our results provide novel sex-dependent targets to prevent renal injury and promote kidney recovery.

## INTRODUCTION

Acute kidney injury (AKI) is a common and serious condition with no specific treatment (1) and worldwide increasing incidence (2). AKI is characterized by a rapid decline of renal function that requires hospital admission and renal function replacement by dialysis if renal failure is severe, leading to high mortality rates (over 50%) (3). Although AKI is reversible and allows at least partial recovery of renal function, repeated AKI episodes increase the risk of subsequent chronic kidney disease (CKD) and cardiovascular disease long after recovery from the original insult (4–8).

Besides infection and toxic drugs, renal ischemia/reperfusion injury (IRI) is a major cause of AKI, which is faced in many clinical situations such as kidney transplantation, partial nephrectomy, renal artery angioplasty, aortic aneurysm surgery and elective urological operations (9). In these conditions, IRI initiates a complex and interrelated sequence of events, resulting in injury and the eventual death of renal cells (9).

Sex differences influence susceptibility, progression and response to AKI. Clinically, an increased mortality rate has been documented among males with acute renal failure (1, 10, 11). In fact, men are more prone to acute and chronic kidney disease and to progress to end-stage renal disease (ERSD) than women, when all-cause incidence rates are considered (12). Studies looking at outcomes in AKI patients have found that sex is an independent predictor of mortality (13–15). Consistent with clinical studies in AKI, animal research has also shown females are protected against renal IRI (16–18). In consequence, pre-clinical studies have been preferentially performed in males although, recently, the importance of defining pathophysiology and disease mechanisms for each sex is increasingly being integrated into biomedical research.

In this study, we have established a renal IRI and recovery model in sexually mature male and female pigs to analyze biochemical markers, histological lesions and molecular changes occurring in pre-ischemic, ischemic and post-ischemic conditions. Thorough analyses of kidney transcriptomic data were performed using Gene Set Enrichment Analysis (GSEA). Besides genes of clinical relevance, gene sets changing their expression pattern in a sex-dependent manner at different time points (basal, injury and recovery) and gene sets differentially expressed at the same time points between males and females were identified. Upon injury, changes in gene set clusters related with immune cell regulation and steroid hormone response, among others, were more prominent in males than females.

Overall, our analysis has brought novel insight into the sex-specific regulation of molecular pathways involved in IRI and recovery. Thus, this study might serve as a resource to better correlate the clinical outcome of IRI with underlying molecular processes that could eventually help to design sex-specific strategies to promote renal regeneration in humans.

## RESULTS

### 1. Renal ischemia/reperfusion injury (IRI) pig model reveals sex-dependent structural and functional renal changes

In order to study the changes in gene expression occurring after ischemia/reperfusion injury (IRI), we proceeded to generate an IRI model in sexually mature pigs. Previous studies on the effects of warm ischemia (19, 20) reported that periods greater than 30 minutes can lead to severe kidney damage and periods longer than 60 minutes irreversible damage. Thereby, ischemic kidney injury was induced in single-kidney female and male pigs by clamping the renal pedicle for 30 minutes and data was obtained before the surgery (PR), five minutes after ischemia (PS) and one week later (WL) (schematized in **Figure 1A**). To confirm our kidney injury–recovery pig model, blood urea nitrogen (BUN) and serum creatinine (SCr) were measured at basal situation before injury (PR), at 5 min post-reperfusion (PS) and at 24 h (1d), 72 h (3d) and one-week (WL) post-reperfusion (**Figure 1B**). SCr and BUN reached their highest values at 24 h post-reperfusion and gradually descended at 3 and 7 days post-surgery for males and females. SCr and BUN levels at WL showed a tendency to be higher than those at PR indicating that the recovery process was still ongoing (**Supplementary Table 1**). No major differences were detected between males and females regarding SCr and BUN levels at any time point (**Figure 1B**). For the scope of this study, we focused on three time points (PR, PS and WL) for subsequent analyses.

**Figure 1.**
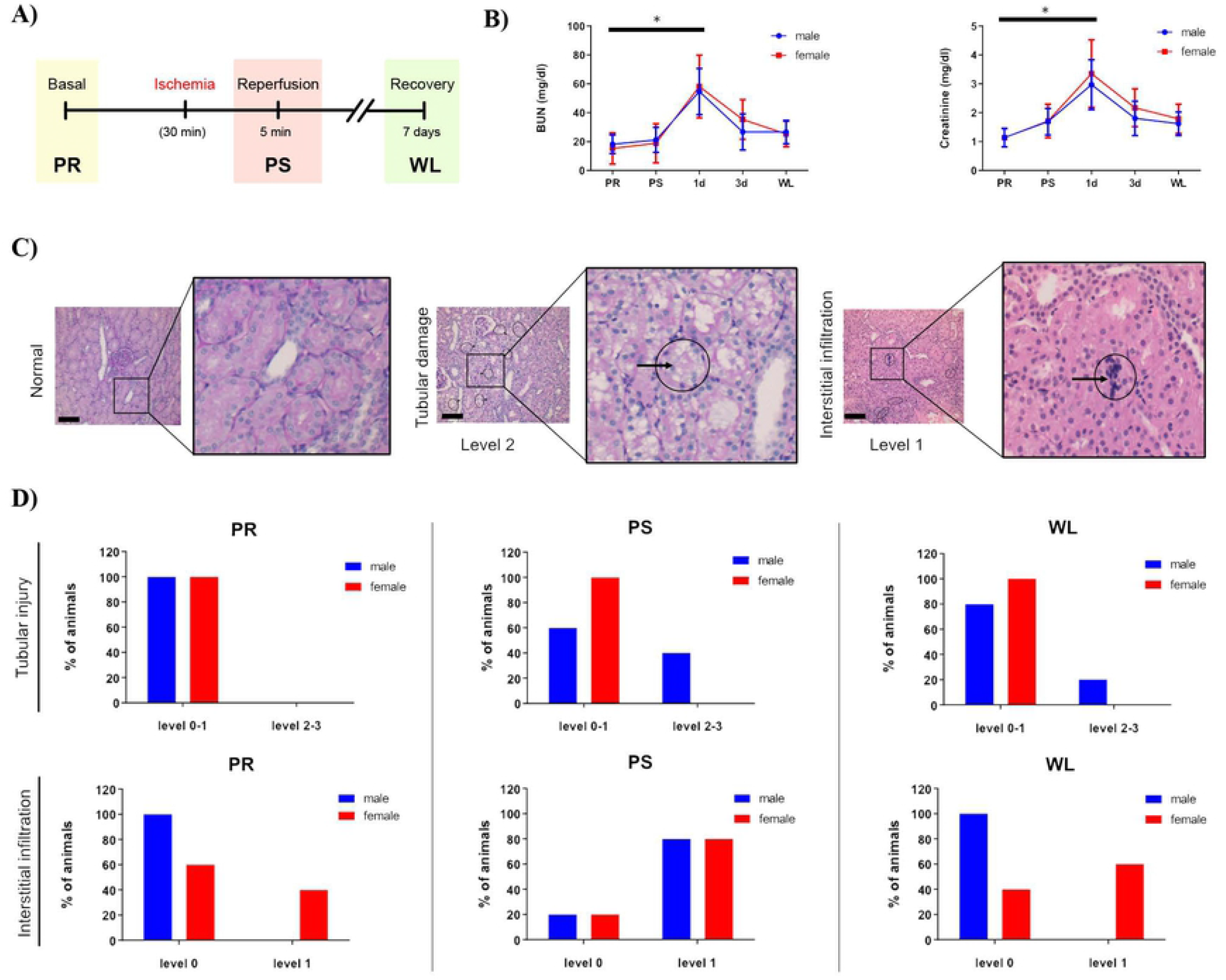
Assessment of biochemical parameters and histological examination following porcine renal ischemia/reperfusion injury. A) Experimental design of renal unilateral IRI following contralateral nephrectomy. Ischemia was induced for 30 minutes. Data were collected before injury, 5 minutes and 7 days following renal clamping. B) Measurement of blood urea nitrogen (BUN) and serum creatinine (SCr) levels in males (blue) and females (red). The “y” axis represents blood urea nitrogen and serum creatinine concentration, respectively, and the “x” axis represents the time points. Average values ± SEM are plotted in the graph (N=5). C) Representative images of different levels of tubular injury and interstitial infiltration in pig kidney. Arrows indicate specifically damaged cells. Magnification = 20X, scale bar = 100 µm. D) Quantification of tubular injury (upper panel) and interstitial infiltration (lower panel) scored by an expert pathologist was classified by group and sex (males in blue, females in red). The y-axis represents the % of animals showing each level of injury or infiltration, respectively. Abbreviations: BUN: blood urea nitrogen PR: pre-ischemia; PS: post-ischemia; WL: one week later. *p<0.05.

Kidney histopathological examination at PR, PS and WL showed near-normal renal morphology with changes associated with sublethal injury including mild interstitial edema and mononuclear infiltration as well as tubular injury associated to brush border diminishment (21–23) (**Figure 1C**). To assess if there was any difference between males and females, we quantified tubular damage (Jablonski scoring system (SS) (24)) and interstitial mononuclear infiltration. Our data revealed that tubular damage was more prominent in males than females (Levels 2-3 of Jablonski SS: 40% of males at PS and 20% at WL vs. 0% of females at PS and WL) (**Figure 1D upper panel**). On the contrary, mononuclear infiltrates (level 1) were more common in females at PR (40% females vs. 0% males), reached similar incidence for both sexes at PS (80% in both males and females) and remained present in 60% of the females but not in males at WL (**Figure 1D lower panel**). Altogether, females present more mononuclear infiltrates than males, both at pre-surgery and at one-week after reperfusion, while males exhibit more tubular injury than females at PS and WL. Importantly, according to biochemical and histological parameters, our data indicate that recovery is still ongoing at 7 days post-reperfusion in both sexes.

### 2. Kidney transcriptome profiles across injury and recovery in female and male pig samples

Next, in order to identify the time- and sex-dependent molecular pathways relevant for IRI and recovery, we performed a microarray-based gene expression analysis using samples from our IRI pig model. To investigate major changes in the transcriptional response before, during and after IRI, we performed a hierarchical clustering of gene expression levels, represented as heatmaps that allowed the identification of common and distinct patterns of regulation between different experimental conditions. In both males and females, time point comparison revealed a similar gene expression pattern between pre-ischemia (PR) and post-ischemia (PS), which was radically different one week later (WL) (**Figure 2A**). Interestingly, sex comparison results indicate that changes in global gene expression observed at PR and PS between males and females disappear during the recovery phase (WL) (**Figure 2B**), with males exhibiting a global female-like phenotype during recovery.

**Figure 2.**
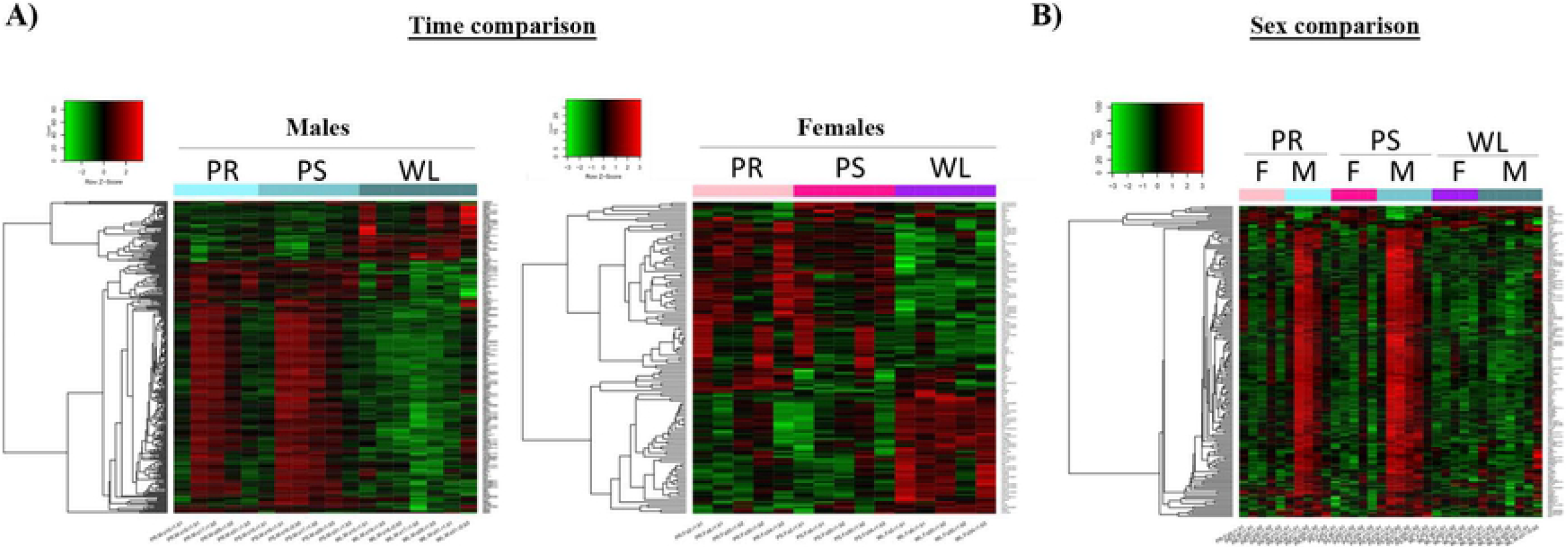
Hierarchical clustering of microarray assays based of kidney porcine throughout renal IRI in a time and sex manner. Gene expression for males and females were compared at different time points (PR, PS, and WL). A) Heatmaps graphically illustrating the differences in the gene expression levels for the time comparison in males (left) and females (right). Similar pattern of expression was observed for both sexes. B) Heatmap representing the difference in expression by comparing males and females at the same time point (sex comparison). The green color represent genes with lower expression and the red color represent the ones with higher expression. Genes represented in the heatmaps have an adj.P. value ≤ 0.25 and |log FC| >=1. Abbreviations: F: female; M: male; PR: pre-ischemia; PS: post-ischemia; WL: one week later.

#### 2a. Validation of microarray data by qRT-PCR

In order to assess the reliability of the microarray data, selected genes changing their expression in a time and sex-dependent manner (*IFIT3, FABP5, CXCl0, CD274, RSAD2*) (**Figure 3A**) were analyzed by qRT-PCR using specific TaqMan probes. Our data show that these genes chosen for the validation presented a similar pattern as in the microarrays, therefore confirming the trustworthiness of the microarray data (**Figure 3B**).

**Figure 3.**
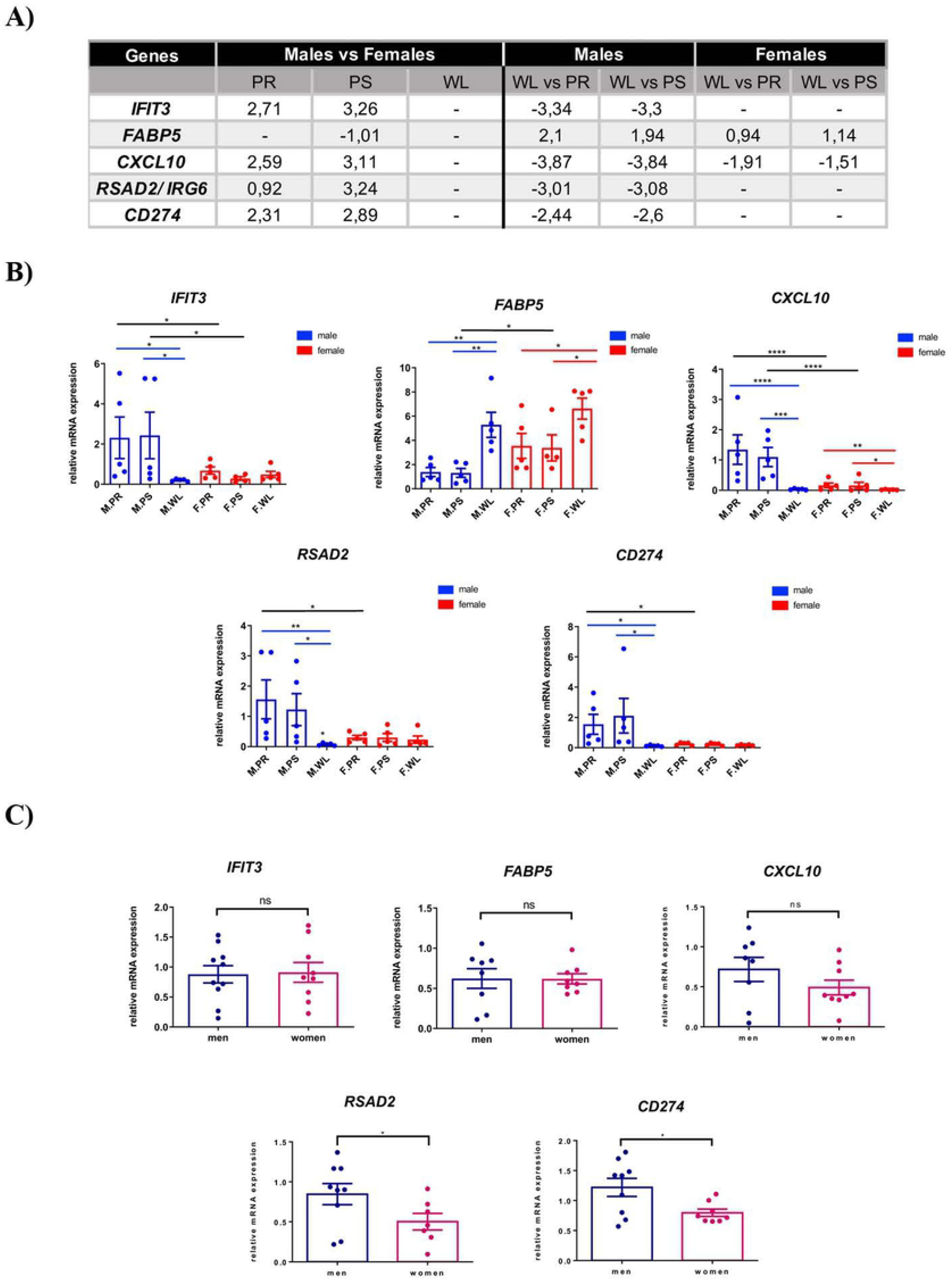
Validation of porcine renal IRI microarray assays by qRT-PCR experiments followed by evaluation of mRNA levels of selected targets in human ischemic kidney biopsies. A) Expression values of five selected targets from microarray assays displaying time and sex differences. Relative mRNA levels of *FABP5, IFIT3, RSAD2, CXCL10, CD274* were measured and compared by qRT-PCR. For the time comparison, the different time points (PR, PS, WL) were compared with each other for each sex (blue male, red female). For the sex comparison, a selected time point in male was compared to the equivalent time point in female. Blue and red lines represent a time comparison in male or female, respectively. Black lines represent sex comparison at equivalent time points. C) RSAD2, CXCL10, CD274, FABP5 and IFIT3 expression levels were evaluated by qPCR in post-surgery (PS) conditions from samples of 36-80 and 53-83 years old men and women, respectively (N>7). *: P-value ≤ 0.05; **: P-value ≤ 0.01; ***: P-value ≤ 0.001; ****: P-value ≤ 0.0001, ns: not significant. Abbreviations: F: female; M: male; PR: pre-ischemia; PS: post-ischemia; WL: one week later.

#### 2b. mRNA levels of pig-selected genes relevant for IRI are conserved in humans

In order to study the conservation of the gene expression pattern in humans, selected genes showing sex-dependent regulation by renal ischemia in pigs (i.e. *IFIT3, FABP5, CXCL0, CD274, RSAD2*) were analyzed in ischemic kidney biopsies from men and women. Briefly, kidney biopsies of normal tissue were obtained from renal cancer patients of both sexes undergoing nephrectomy. Non-tumoral post-ischemic tissues were collected after 30 minutes of ischemia, approximately, thus corresponding to the post-surgery (PS) condition in our pig model. Next, we tested the mRNA levels of the selected genes in these post-ischemic kidneys of men and women by quantitative PCR (qRT-PCR). Importantly, from the five genes analyzed, *RSAD2, CXCL10* and *CD274* showed the same expression pattern observed in the pig model (*RSAD2*: men: 0.8462 ± 0.1322, women: 0.5015 ± 0.1036; *CXCL10*: men: 0.7169 ± 0.1500, women: 0.4909 ± 0.0917; *CD274:* men: 1.219 ± 0.1505, women: 0.7965 ± 0.0608), while no differences were detected for *FABP5* or *IFIT3* between sexes (**Figure 3C**). Overall, a partial correlation was found between the mRNA levels of both species, under ischemic conditions.

Altogether, our data suggest that *RSAD2, CXCL10* and *CD274* might serve as noninvasive surrogated biomarkers to predict ischemic injury and recovery in human kidneys.

#### 2c. Time comparison reveals differentially expressed genes throughout renal IRI

We have analyzed those transcripts altered throughout IRI (up- and down-regulated) to identify common and exclusive genes for each time point (PR, PS, WL). In males, 174 genes were conserved between the WL vs. PR and WL vs. PS comparisons, 52 genes were exclusive of WL vs. PS comparison and 50 genes were only found in the WL vs. PR comparison. From the eight genes that are different between PS and PR, only two (*LEAP2, MIR505*) are exclusive of this comparison (**Figure 4A**). Our results revealed similar patterns for each comparison in females, although lesser genes were altered compared to males. Specifically, 59 genes were conserved between the WL vs. PR and WL vs. PS comparisons, 19 genes were exclusive of WL vs. PS comparison and 37 genes were only found in the WL vs. PR comparison (**Figure 4B**). Interestingly, one gene (*FOS*) was shared between the WL vs. PR and the PS vs. PR comparisons. Finally, from the eight genes differentially expressed between PS and PR, four are specific for this comparison (*NKL, KLF5, DNAJB1* and *CPE*).

**Figure 4.**
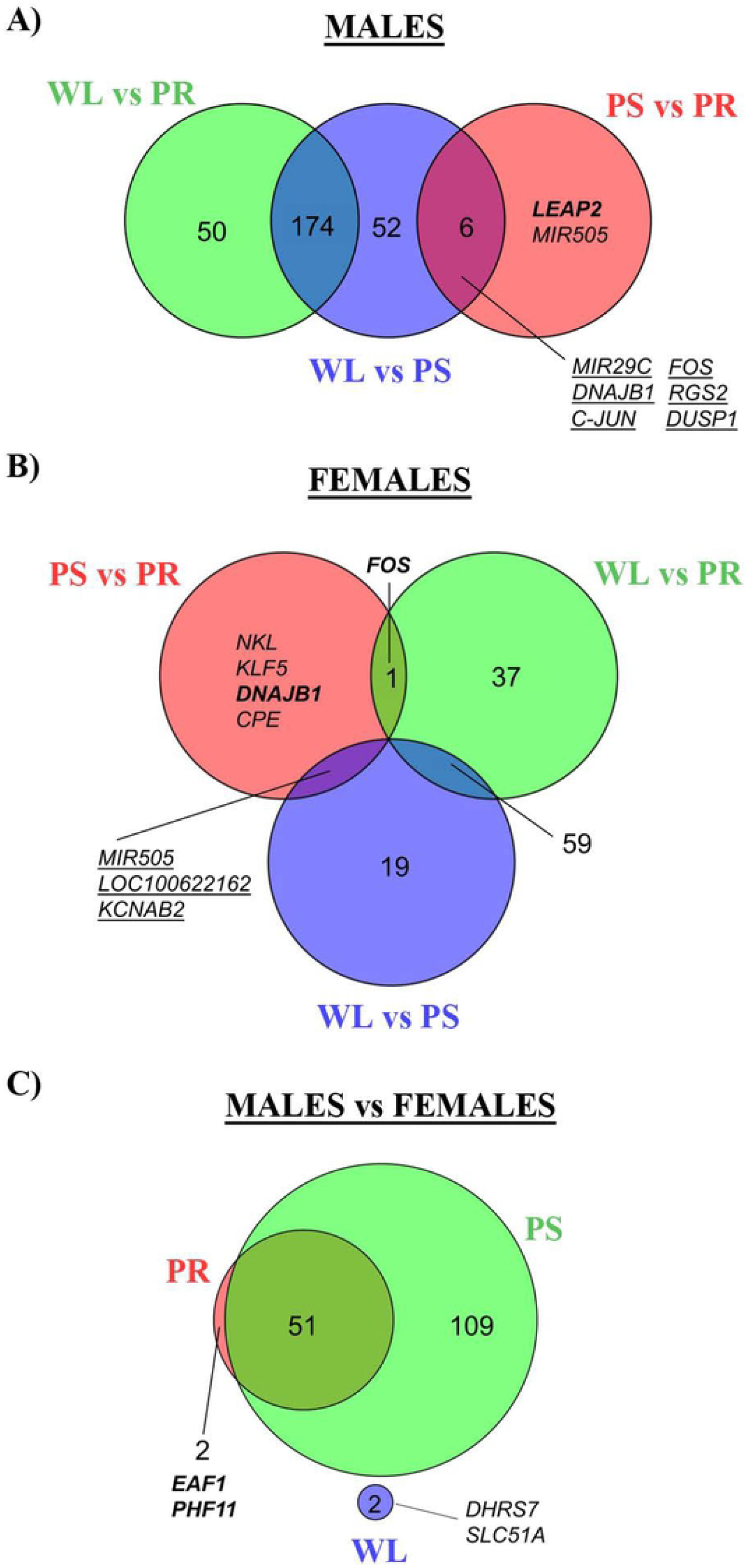
Venn diagrams of time and sex comparisons between males and females throughout renal IRI. Venn diagrams depicting the number of commonly regulated genes A) in male and B) in females at different time point comparisons: PS vs. PR, WL vs. PS and WL vs. PR. Males showed an overall higher number of regulated genes. C) Venn diagrams depicting the number of commonly regulated genes in the sex comparison at different time points: PR, PS and WL. Only two genes were commonly regulated one week following injury. Genes represented only in italic are down-regulated, whereas genes in bold and italic are up-regulated. Genes underlined are both up- and down-regulated in respective comparisons. Complete gene tables are available in supplemental material. adj. p-value ≤0.25 and log |FC |≥ 1. Abbreviations: F: female; M: male; PR: pre-ischemia; PS: post-ischemia; WL: one week later.

Altogether, our data suggest that renal injury and recovery processes have a lower impact in females than males. The overall number of differentially expressed genes in renal IRI and recovery in male and female pig kidneys are reported in the supplementary figure 1 (adj. p-value ≤0.25, without considering the fold change, **supplementary figure 1**).

#### 2d. Sex comparison of gene expression during renal injury and repair

Fifty-one of the 53 differentially expressed genes between sexes at basal (PR) remained unchanged after injury (PS), while 109 genes changed in males during this phase (**Figure 4C**). It is very relevant to state that all gene expression changes found between males and females at PR and PS disappeared at WL and only two genes, *SLC51A* and *DHRS7*, were differentially expressed between sexes in this phase. The number of differentially expressed genes throughout renal IRI and recovery between males and females at the same time point are reported in supplementary figure 2 (adj. p-value ≤0.25 without considering the fold change, **supplementary figure 2**).

### 3. Role of androgens in the regulation of differentially expressed genes (DEG) during IRI

Our results showed a clear difference in gene expression between males and females during IRI and recovery process, thus we postulated that sexual hormones might have a role. To better assess it, we compared the gene expression profile of the top DEG between male and female pigs over IRI with data from a single castrated male (CM) using the Ingenuity Pathways Analysis (IPA) software. Interestingly, the CM sample, which was not included in the enrichment pathway analyses, phenocopies the gene expression pattern of females at basal conditions (PR) (**Figure 5A**). Moreover, the CM does not completely follow the gene expression pattern of males at PR and WL (**Figure 5B**). Specifically, genes that follow a putative androgen-dependent expression in PR (i.e. *UBD, IFIT3, CXCL11, FBG, FGG, MX1, IFIT1* and *CXCL10*) show an opposite direction at WL between males and CM. On the other hand, those that are common between males and CM (i.e. *CKAP2, CENPF, CDC20, KIF20A, CCNA2, EPHB3, C6, SLC6A19* and *FABP5)* in PR, retain the same pattern of expression at WL. These results suggest that androgens and male sexual hormones may contribute to the sexual dimorphic expression pattern observed at basal conditions, in renal injury and during recovery.

**Figure 5.**
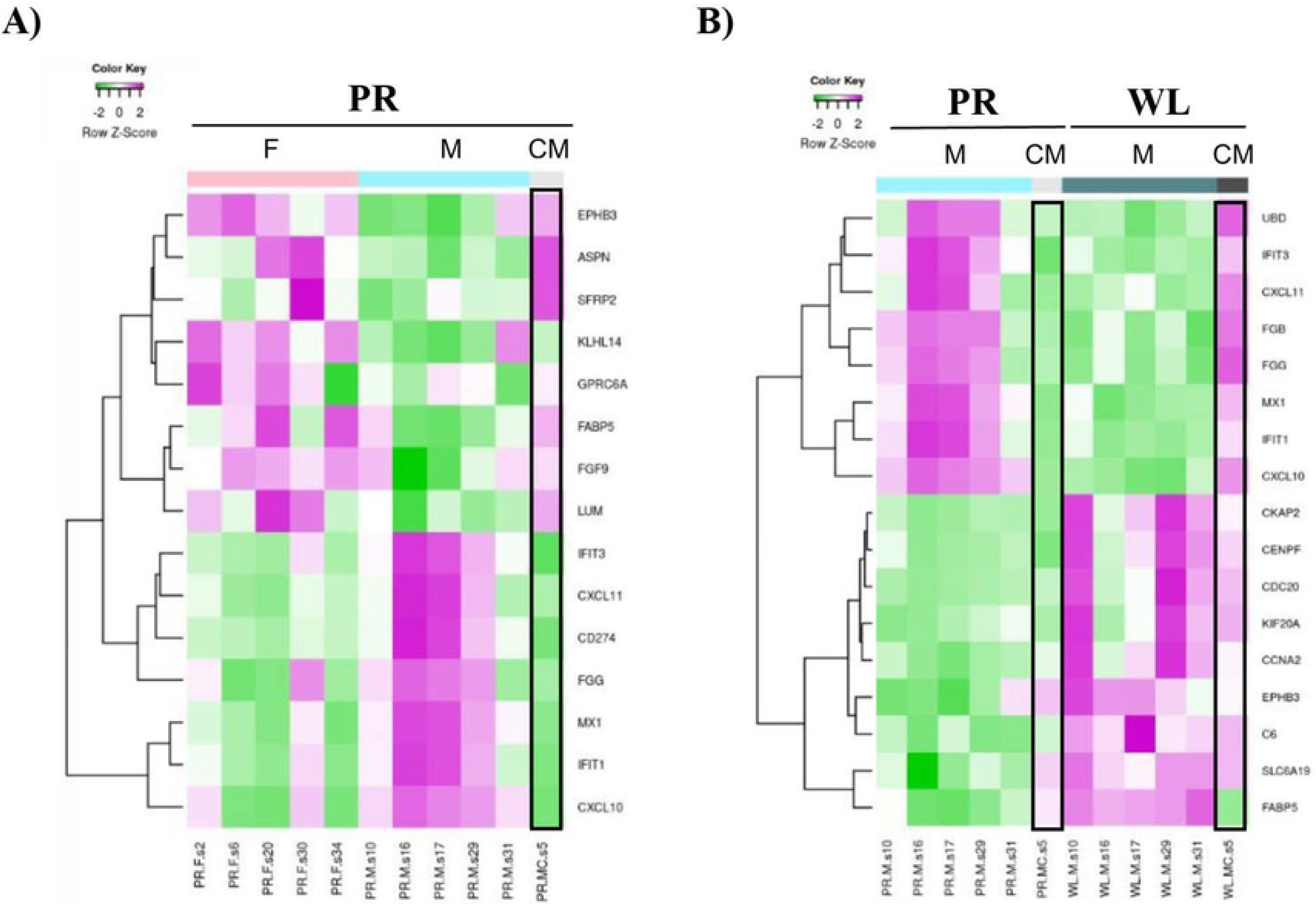
IPA heatmap gene expression representation of top regulated genes. Microarray data files of pig experiments were uploaded in IPA software. Results were reported in hierarchical clustering of top up and down regulated genes in A) a sex-(MPR vs. FPR) and B) time-(MPR vs. MWL) comparisons. Data from a castrated male was compared with male and female pig expression patterns. The castrated male showed a gene expression pattern similar to females. Abbreviations: F: female; M: male; CM: castrated male; PR: pre-ischemia; WL: one week later.

### 4. Gene Set Enrichment Analysis (GSEA) reveals the importance of immune system related pathways during the recovery process in males

Our study has identified differentially expressed genes (DEG) between female and male pig kidneys at basal conditions (PR), injury (PS) and recovery (WL). To gain mechanistic insight into time- and sex-related differences that govern renal injury and recovery processes, we aimed to identify which biological pathways are differentially expressed between groups using Gene Set Enrichment Analysis (GSEA).

As an example of the GSEA, here we show the comparison between males and females at one-week post reperfusion (M.WL vs. F.WL). For this particular comparison, the GSEA showed that over-represented clusters (in red) in males include: regulation and production of cytokines and interleukins, immune somatic recombination, microtubule cytoskeleton organization, actin organization and mitotic cycle transition. On the other hand, under-represented clusters (in blue) contain nodes related with metabolism of fatty acids and steroid hormones, nucleotide biosynthetic processes, amino acid catabolism and response to xenobiotic stimulus, amongst others (**Figure 6A**). Moreover, deeper analysis of each of these nodes led to specific gene sets. For instance, the <u>somatic recombination immune node</u> (up-regulated) includes gene sets like lymphocyte activation or B-cell differentiation; while the down-regulated <u>node of fatty acids and steroid hormones</u> includes metabolism of steroids or metabolism of lipids (**Figure 6B**). We performed the same analysis for the other sex and time comparison before, after injury and during recovery. The lists of 10 top up- and down-regulated gene sets enriched for each comparison are reported in the **supplementary Tables 2-10**. Overall, our results revealed the sex-specific regulation of gene sets upon IRI.

**Figure 6.**
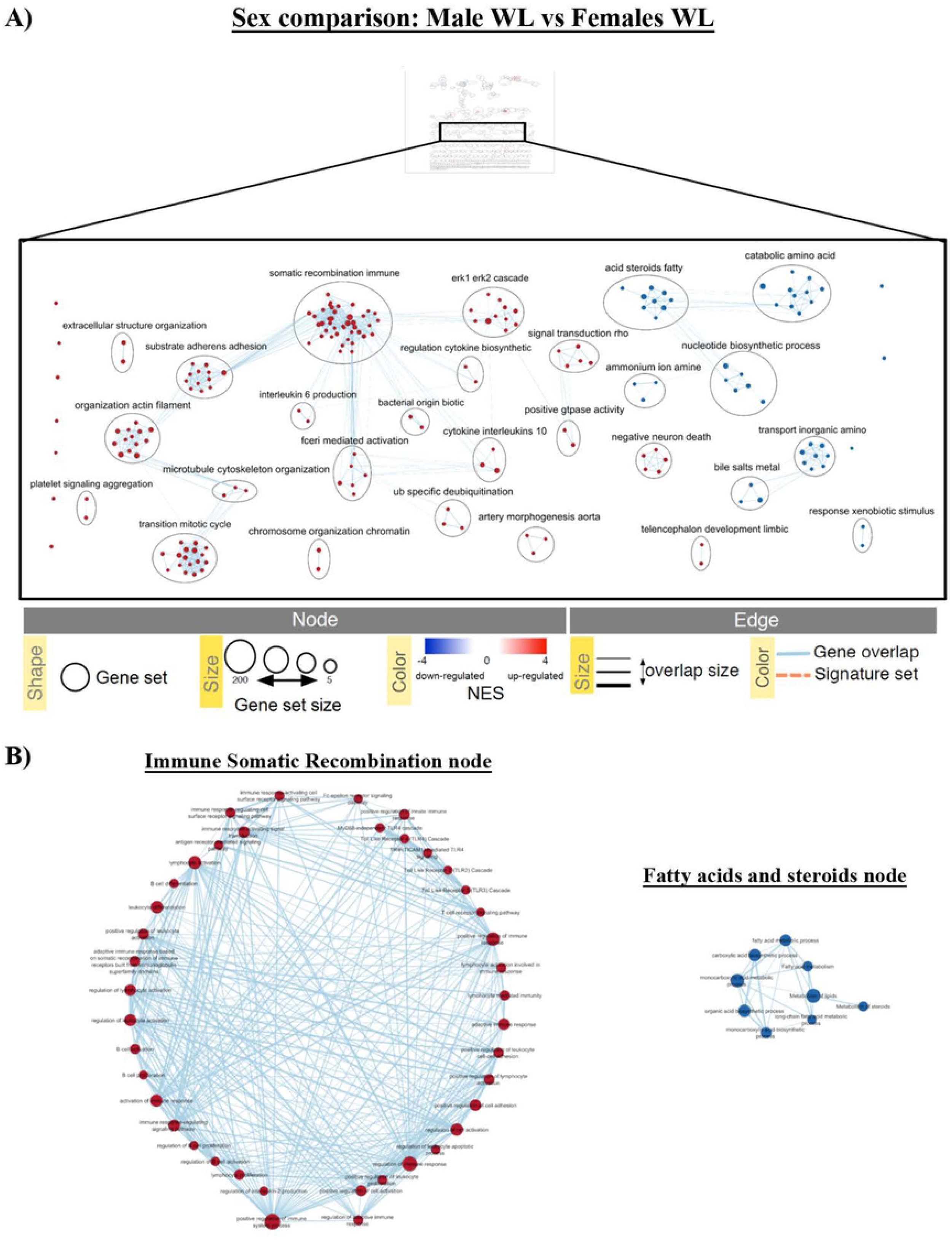
Enrichment map example of over-represented genes in individual sex comparisons M.WL vs. F.WL following GSEA analyses. A) Representation of different clusters (nodes) regulated in the comparison. The map allows visualization of clusters containing nodes in which red and blue represent up- or down-regulated gene sets for each node, respectively. The clusters take their name from the most common containing names of the nodes within the cluster. B) Example of the different gene sets that form somatic recombination immune and acid steroids fatty nodes, where red and blue nodes represent up- or down-regulated gene sets, respectively. (FDR: 0.01-0.1).

### 5. Grouped enrichment analyses of sex and time comparisons show the gene sets controlling renal injury and recovery

Besides individual comparisons and to get insight into the overall processes, we performed grouped comparison visualization of the enrichment analyses. Heatmaps including the six time comparisons (F.PS vs. F.PR; FWL vs. F.PS; F.WL vs. F.PR; M.PS vs. M.PR; M.WL vs. M.PS; M.WL vs. M.PR) or the 3 sex comparisons (M.PR vs. F.PR, M.PS vs. F.PS, M.WL vs. F.WL) were created. The addition of multiple comparisons in a single enrichment map hindered an effective visualization of gene sets involved in chosen clusters. Nine clusters containing high numbers of gene sets, which revealed their prominence in the processes under study, were selected from the enrichment maps. The NES (normalized enrichment score) value of each of the gene sets included in the nine clusters were indicated in heatmaps. The nine clusters are: (I) immune cell regulation, (II) morphogenesis development migration, (III) ion transport transmembrane, (IV) apoptotic intrinsic extrinsic, (V) oxygen levels hypoxia, (VI) alcohol biosynthetic process, (VII) steroid hormone response, (VIII) regulation hormone secretion and (IX) phagocytosis endocytosis and invagination. Interestingly, the regulation of gene sets of a distinct cluster varies amongst different comparisons (examples in **Figure 7A** and **Figure 8A**).

**Figure 7.**
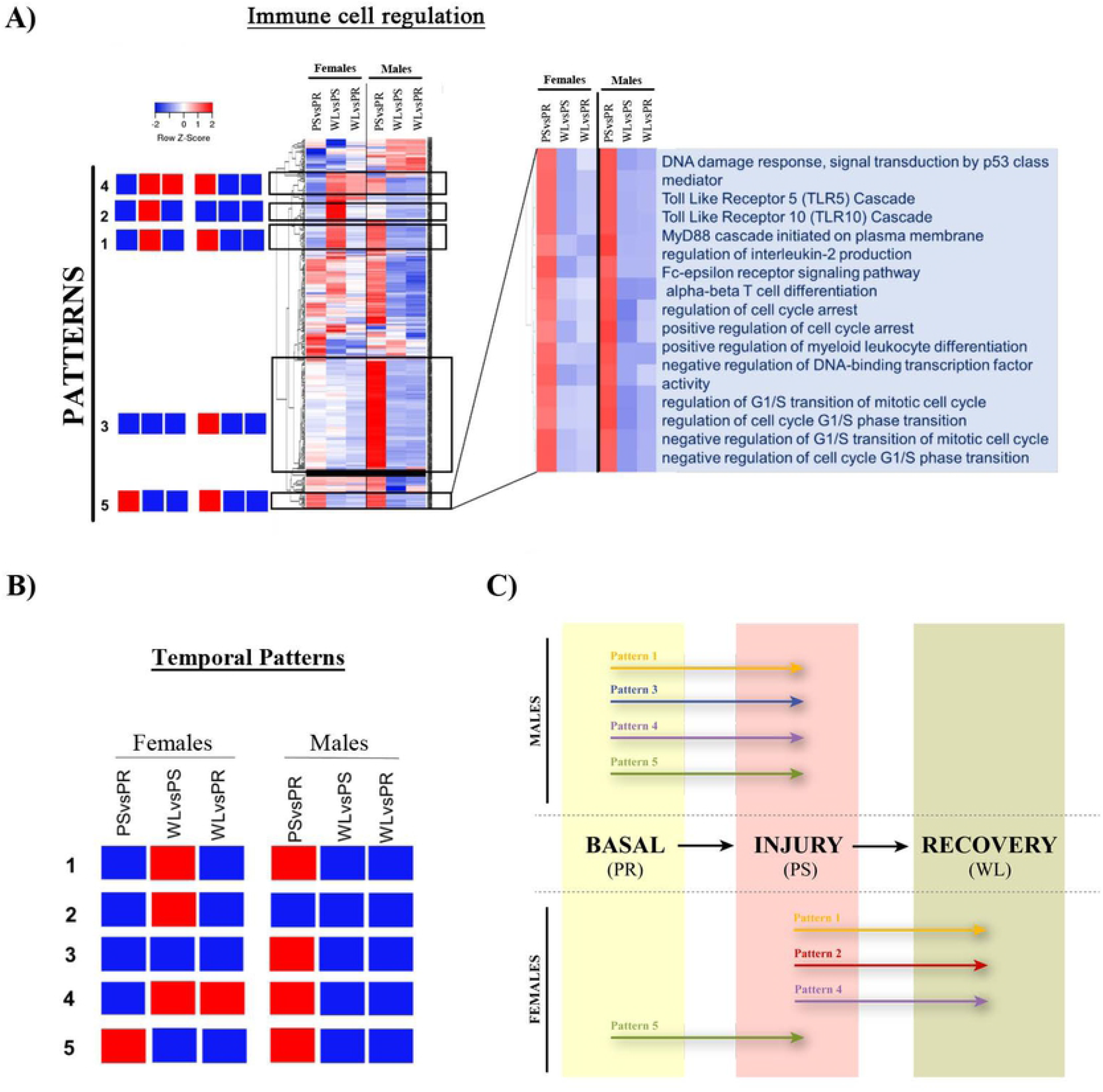
Patterns of gene sets regulation in male and female kidneys throughout renal IRI in the time comparison. An example of the gene sets of selected clusters represented by hierarchical clustering. A) Heatmap (of time comparisons) was created with the normalized enrichment score (NES) values of the gene sets calculated by GSEA analysis. The red and blue colors refer to gene sets that are over- or under-represented in the heat-maps. B) Five prominent patterns for time comparison were determined. C) A summary of these five temporal patterns is depicted in a diagram, where patterns displayed in each sex are illustrated by a colored arrow positioned at the time point where they are up-regulated (PR, PS, WL). Abbreviations: PR: pre-ischemia; PS: post-ischemia; WL: one week later.

**Figure 8.**
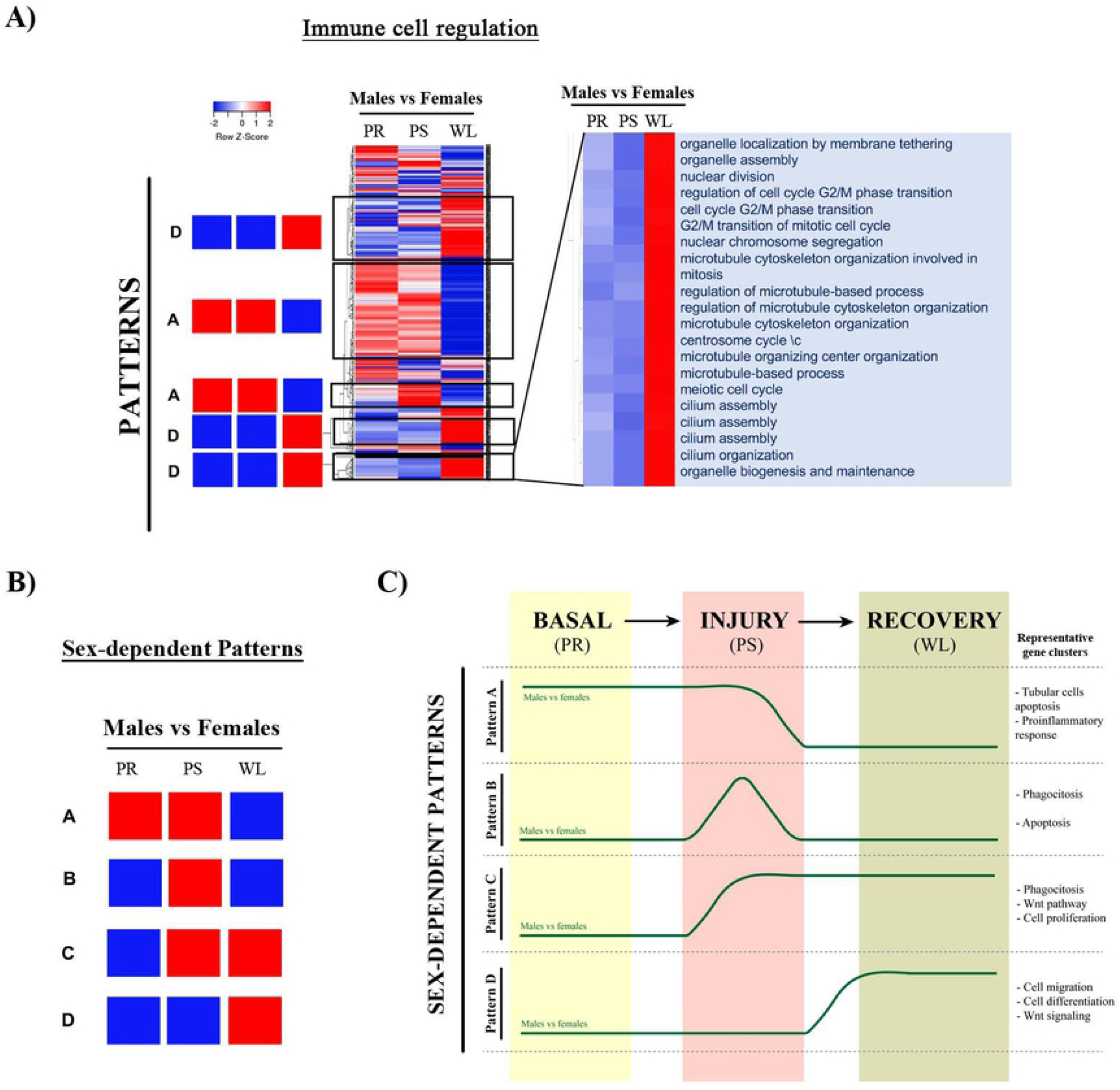
Patterns of gene sets regulation in male and female kidneys throughout renal IRI in the sex comparison. Gene sets of selected clusters were represented by hierarchical clustering. A) Heatmaps (of sex comparisons) were created with the normalized enrichment score (NES) values of the gene sets calculated by GSEA analysis. The red and blue colors refer to gene sets that are over- or under-represented in the heat-maps. B) Four prominent patterns for sex comparison were determined. C) A summary of these four temporal patterns is depicted in a diagram. Abbreviations: PR: pre-ischemia; PS: post-ischemia; WL: one week later).

#### 5a. Grouped GSEA analyses reveal different temporal gene set regulation patterns after renal IRI and recovery

Heatmaps representing gene sets of previously selected clusters (for instance, the Immune cell regulation cluster shown in **Figure 7A**) allowed the visualization of five prominent temporal patterns of expression that are schematically represented in **Figure 7B**. To simplify our analysis, we focused on gene sets that are over-regulated, assuming that this leads to higher activity of those genes involved in IRI events. Importantly, gene sets from the nine clusters can follow different or similar temporal patterns revealing coordinated expression (**Figure 7B** and **tables 1 to 5**). The five temporal patterns that we have identified are (**Figure 7C**):

1. **<u>Pattern 1</u>** includes gene set clusters that are over-represented during the recovery process in females (WL vs. PS) and in the injury process in males (PS vs. PR) (see **Table 1** for complete gene sets included in pattern 1).
2. **<u>Pattern 2</u>** is composed of pathways over-represented during the recovery process (WL vs. PS) in females but never found in males (**Table 2**).
3. **<u>Pattern 3</u>** includes gene sets that are only over-represented in males during injury (PS vs. PR) (**Table 3**).
4. **<u>Pattern 4</u>** is composed of gene sets over-represented during the recovery process (WL vs. PS) and at one-week post-reperfusion (WL vs. PR) in females; and also over-represented only during injury in males (PS vs. PR) (**Table 4**).
5. **<u>Pattern 5</u>** involves pathways that are over-represented only during injury in both sexes (PS vs. PR) (**Table 5**).

#### 5b. GSEA analyses reveal four sex-dependent gene set regulation patterns

Finally, we have created heatmaps for the sex comparison (for example, for the immune cell regulation cluster shown in **Figure 8A**), which revealed four prominent sex-dependent patterns (from A to D) schematically shown in **Figure 8B**. We performed the same type of analysis as for time comparison to regroup the gene sets and clusters that shared similar expression pattern (**Tables 6 to 9**) (**Figure 8C**).

- **<u>Pattern A</u>** includes gene sets that are up-regulated in males vs. females both at basal conditions (PR) and after injury (PS) (**Table 6**).
- **<u>Pattern B</u>** includes over-represented gene sets in males at injury (PS) (**Table 7**).
- **<u>Pattern C</u>** includes pathways over-represented in males at injury (PS) and after one-week reperfusion (WL) (**Table 8**).
- **<u>Pattern D</u>** is followed by pathways over-activated in males one week after reperfusion (**Table 9**).

## DISCUSSION

Ischemia is the most common etiology for acute kidney injury (AKI) and one of the main contributors to morbidity and mortality in the hospital setting, as it affects 1 out of 5 patients in emergency admissions (25). Experimental studies have also shown that AKI is associated with mild-to-moderate acute injury in organs distant form the kidney such as the liver, lung or brain therefore precipitating or aggravating other conditions that may have significant impact on patients’ morbidity and life expectancy (26). The initiating insult might be irreversible but, in many cases, timely intervention to restore renal perfusion may mitigate the severity of evolving ischemic AKI, by preventing still functioning tissue from progressing to overt injury. AKI occurrence also displays sex differences, men being generally more prone to suffer from AKI, to progress more frequently to chronic kidney disease (CKD) and to end stage-renal disease (ESRD) (27).

### Novel renal Ischemia/Reperfusion Injury (IRI) pig model

Pigs represent the gold standard model for renal transplantation research and studies involving IRI in tubule interstitial fibrosis development. Pigs present advantages over other animal models because their similarities with humans (e.g. genome, size, metabolism and renal anatomy) (28, 29), being some biochemical parameters identical (e.g. Scr and BUN) (24, 25). Importantly, the size of their kidney allows sample collection at different time points from same animal, overcoming the individual variability and disparity that might occur in rodents.

Our IRI pig model presents the highest SCr and BUN values, markers of renal injury, at 24 hours post-reperfusion in both males and females. Their levels gradually descend and remain slightly elevated seven days after injury, which indicates an ongoing recovery process. This reproduces the course of ischemic AKI observed in patients, where the process from insult to first evidence of recovery takes between 7 and 21 days (25). Although both sexes showed similar levels of SCr and BUN, kidney histopathological examination revealed sublethal injury with higher mononuclear infiltrates in females than males. These data correlate with the immune response sexual dimorphic pattern observed in humans (31). On the other hand, tubular injury associated to brush border diminishment was still present in males at 7 days post-injury. This indicates that the renal recovery after IRI is delayed in males compared to females, suggesting a role for sexual hormones in this process.

### Identification of sexual dimorphism in the time-specific gene expression controlling renal IRI and recovery

The first key result of this IRI pig model is that males exhibit stronger global gene expression changes during injury and recovery than females. It is striking the opposite expression pattern observed in males compared to females even at basal (PR) or during injury (PS), which is completely reversed with males acquiring a female-like gene expression pattern at 7 days post-surgery (WL). Altogether, these data point to a clear role for the sexual hormones in the protection against IRI and recovery after renal injury.

Only two genes (*SLC51A* and *DHRS7*) remain differentially expressed at 7-day post-surgery between males and females. *DHRS7* encodes for the seventh member of the short-chain dehydrogenases/reductases (SDR) family, which metabolize many different compounds, including steroid hormones (32). *SLC51A* encodes the alpha subunit of the organic solute transporter alpha/beta (OSTα/β), which is a heteromeric solute carrier protein that transports bile acids, steroid metabolites and drugs into and out of cells (33). The differential regulation between sexes of genes that metabolize and transport sex steroid hormones during recovery suggests a link between their expression and renal repair after injury.

Amongst the genes with differential expression between females and males at basal conditions or during injury, but exhibiting similar levels during recovery, the ones with the highest differences are those related with immune and inflammatory processes. For example, interferon signaling pathways related genes (*MX1, IFIT3* and *GBP1)*, interferon responding genes (*CXCL9, CXCL10* and *CXCL11*), programmed cell death 1 ligand 1 (*PDL1*/*CD274*), inflammatory response protein 6 (*IRG6*/ *RSAD2*) and a protease that cleaves complement components *C2* and *C4* (*MASP2*) (34). All these genes are strongly and significantly overexpressed in males compared to females at basal conditions or right after injury, with no differences at seven-days post-injury indicating that they acquire a female-like expression phenotype.

Nevertheless, albeit presenting similar expression of immune related genes at recovery, the histopathological analysis shows higher mononuclear infiltrate in females. A possible explanation is that same ligands can have different effects depending on the cell type. For instance, chemokines CXCL9, CXCL10 and CXCL11 are ligands of the CXCR3 receptor and play important roles in the activation and stimulation of the immune system against foreign antigens (35). However, CXCR3 positive T regulatory (Treg) cells infiltration are beneficial for proper kidney allograft function (36). This dual effect could explain why females are more protected against IRI than males. One of the limitations of our study has been the poor performance of available antibodies to detect pig proteins by WB and IHQ assays, thus enabling us to correlate differential gene expression with immune cell infiltrates in kidney tissues.

### Role of sexual hormones in the regulation of IRI and renal recovery controlling genes

Our data showed that the expression pattern of putative sex-regulated genes in the castrated male was closer to females than males, confirming the impact of male sexual hormones on IRI and recovery. This is the case, for example, for *FABP5, CD274, IFIT3* and *CXCL10* genes, likely indicating the androgen-dependent regulation of their expression. In fact, *FABP5* has been found to be a potential therapeutic target in prostate cancer, an androgen-dependent cancer type (37). Moreover, androgens have been shown to up-regulate *CXCL10* expression in prostate epithelial cells (38).

### Comparison between humans and pigs data: PDL1 as a candidate

An important part of our study was to prove the correlation between pigs and humans, so our discoveries could be used to treat renal IRI. Our data from human samples show that the expression levels of *CXCL10, RSAD2* and *CD274* (PDL1) are lower in females than males, similar to what was observed in our pig model. Amongst them, CD274/PDL1 is one of the most interesting candidates. This protein is a ligand of PD-1, a negative co-stimulatory molecule expressed by T lymphocytes, monocytes, dendritic cells, and B cells (39). The interaction between PD-1 and PDL1, present on antigen-presenting cells and tumor cells, constitutes an immune checkpoint through which tumors can induce T-cell tolerance and avoid immune destruction (39). It appears that PDL1 on non-immune cells participates in Treg-mediated protection against kidney IRI and AKI (40). However, further research is required to study how PDL1 lower levels in females can protect them against injury.

### Time and sex-dependent IRI and repair pathways

Besides individual genes, our –omics data allowed the identification of clusters containing gene sets relevant for processes associated with renal IRI and recovery, providing evidence of their sex- and temporal-regulated fashion.

#### a) Sex and time comparison of gene sets of most prominent clusters

Here we have identified **five temporal patterns** for different gene clusters and **four sex-dependent patterns**. The behavior of these temporal and sex-dependent patterns is summarized in **Figure 7C and Figure 8C**. First, early after IRI (PR to PS), genes following <u>pattern 5</u> are activated in both males and females, but males also specifically activate the genes following <u>pattern 3</u>. Interestingly, <u>pattern 1</u> and <u>pattern 4</u> include genes expressed after injury in males, but one week later (recovery) in females. Finally, the <u>pattern 2</u> is specific for females one week after the injury. In order to understand the meaning of this temporal and sex-dependent regulation, as well as the clusters activated at each time point, we propose the following:

In the <u>early phases of renal IRI</u>, reduced oxygen supply to metabolically active tubular epithelial cells lowers oxidative metabolism and depletes cell supplies of high-energy phosphate compounds. Reperfusion restores the oxygen supply, which results in mitochondrial impairment, enhances oxygen free radicals formation and, therefore, causes more injury (41). Interestingly, males and females react differently to this situation. Males activate gene sets in response to a decrease in oxygen levels and hypoxia (**Pattern 3**), while females show negative regulation of angiogenesis during the recovery phase (**Pattern 2**). Altogether, our data likely indicate that males suffer more from the lack of oxygen than females.

<u>Tubular epithelial cell apoptosis</u> is the key pathophysiological alteration occurring in IRI, and defines the extent of the damage to kidney function (41). This process mainly occurs through the intrinsic pathway (42) by p53, which is highly activated in males at both basal and injury conditions (**Pattern A**), suggesting that males are preferentially affected by apoptotic damage. This fits well with the histopathological results showing more tubular injury associated to brush border diminishment in males than females. Androgens are known to inhibit apoptosis and promote growth (43, 44). However, upon cellular stress, they can also promote stress-mediated apoptosis by enhancing mitochondrial translocation of the proapoptotic protein Bax, which plays a critical role in the intrinsic apoptotic pathway via mitochondrial membrane permeabilization (45).

Another process leading to tubular epithelial cell death is <u>necrosis</u>. Necrotic cell death is accompanied by the release of immunogenic cellular components collectively known as damage-associated molecular patterns (DAMPs), which cause severe tissue damage, leading to systemic inflammation (46). Apoptosis and regulated necrosis can occur at the same time in the same kidney compartment, as they are not mutually exclusive and coexist in many renal pathological conditions (41). Gene sets related with necrotic cell death are upregulated in females during recovery (**Pattern 2**), occurring later than apoptosis in males. Interestingly, our results point to sex hormones as a relevant factor pushing towards apoptosis or necrosis in front of the same trigger and intensity event.

<u>During IRI</u>, both sexes activate gene sets related with DNA damage, the innate immune response, T cell activation, cytokine secretion and cell cycle arrest, which are downregulated during recovery (**Pattern 5**). Together with tubular epithelial cells, macrophages produce proinflammatory cytokines, thus contributing to injury. Gene sets and clusters controlling these pathways are preferentially upregulated in males (**Pattern 3**). Besides, males also present enhanced activation of pro-inflammatory pathways (e.g. *TNF alpha* and *IFNγ* production, *NFKB* signaling and complement cascade) not only during injury but also at basal situation (**Pattern A**). Concomitantly, negative regulation of MAP kinase activity and positive regulation of hormone metabolism processes also occur in both sexes, contributing to the reestablishment of cellular homeostasis.

Although the immune response has an important role during these processes for both sexes, gene sets involved on immune cell regulation, mononuclear cell migration, leukocyte chemotaxis, phagocytosis and engulfment are activated at different time points in males and females. Genes following patterns 1 and 4 are active at injury in males but enhanced in females during recovery, which correlates with the apoptotic and necrotic events occurring in males and females at these time points, respectively. Our histology data also revealed higher mononuclear infiltrates in females at recovery, which is in agreement with gene sets following **pattern 2** (upregulated in females during recovery) controlling inflammatory response, TNF alpha production, humoral response and adaptive immune responses. These processes are even clearer when we compare male and female at the same time points, which reveals that genes related to phagocytosis are more activated in males at injury and during recovery (**Pattern C and Pattern D**).

The renal tubular epithelium has a huge capacity for <u>regeneration after injury</u>. During the repair process, surviving tubular cells actively proliferate and differentiate into mature tubular cells to reconstruct their functional structures. Regeneration of the tubular system is essential for recovery from AKI and a clear marker of patient morbidity (41). The clinical end-point of abnormal repair is chronic kidney disease that is reflected, histologically, by tubular atrophy and renal fibrosis due to myofibroblast proliferation and deposition of extra-cellular matrix (25). Regeneration involves actions of endogenous inhibitors of inflammation, up-regulation of repair genes, actions of the immune system, clearance of necrotic and apoptotic cells and tubular regeneration (25). Gene sets regulating these pro-regenerative processes through the immune system are activated one week after the injury in our model, especially in females (**Pattern 4**). We also observed that gene sets related to extra-cellular matrix and cellular migration are upregulated in males during injury (**Pattern 3**), which indicates an effort to replace lost cells to repair the tubular system, a phenomenon that usually occurs within less than a week (41).

As shown by gene sets included in **Pattern C**, males show a positive regulation of Wnt signaling pathway, endocytosis and engulfment at injury and during recovery, likely reflecting that injury has a stronger effect in males. It is also apparent a negative regulation of the intrinsic apoptotic pathway during injury (**Pattern B**) possibly due to the induction of EMT (epithelial-mesenchymal transition) in males (**Pattern 3**). The EMT activation together with increased epithelial and endothelial cell proliferation that occurs in males during injury and recovery (**Pattern C**) might allow males to recover from the ischemic insult. Moreover, the migratory capacity provided by EMT enables these transitional cells to invade the basement membrane and repopulate the injured tubules (47). **Pattern D** shows that males at recovery exhibit augmented transcriptional programs related with endothelial cell migration and differentiation, kidney development, morphogenesis and epithelium recovery, associated to the activation of canonical and non-canonical Wnt signaling. Activation of Wnt/β-catenin seems to be instrumental for tubular repair and regeneration after AKI, recapitulating the role of Wnt signaling in kidney embryonic development (48).

## Conclusions

Our results show that sex hormone have an impact on the type of gene sets regulated in the kidney during IRI and recovery and also on the timing of their activity. Steroid biosynthesis, hormone secretion and hormones transport are up-regulated in males compared to females in basal conditions and after injury. These differences are abolished one-week after injury, fitting with the feminized gene expression pattern shown by males during recovery, which might likely represent a survival mechanism to diminish androgen promotion of stress-mediated apoptosis. Altogether, our study provides a template to further characterize renal IRI in a temporal and sex specific manner that might bring us one step closer to the development of effective treatment strategies for kidney diseases in the human population.

## MATERIALS AND METHODS

### Animals

This study was conducted using farm pigs, hybrids between Large White and Landrace. Five females, five males and one castrated male of four months old, free of specific pathogens, between 30–40 kg of weight were included in this study. This age range was chosen due to the sexual maturity of the animal, allowing hormone effects. All animal care and procedures were performed in accordance with the requirements of the European laws on the protection of animals used for scientific and experimental purposes (86/609 EEC), and has the approval by the Experimental Ethics Committee of the Vall d’Hebron Institute of Research (VHIR) (34/08 EAEC).

### Experimental design

On the first surgical day all pigs underwent left nephrectomy. On the second day (one week later), the right kidneys were subjected to 30 minutes of warm arterial ischemia followed by 5 minutes of reperfusion and allowed for seven days of recovery (**Figure 1A**). Overall, three kidney biopsies were collected for each animal: prior to injury (PR), 5 minutes following 30 minutes of ischemia (PS) and one week after ischemia (WL). In addition to tissues, blood samples were collected at the different time points of the experiment including 1 and 3 days following ischemia.

### Assessment of renal injury by serum analysis

Creatinine and urea serum levels were analyzed on blood samples obtained through the cannulation of the carotid artery and the internal jugular vein (placed during the first surgery). The catheters subcutaneously tunneled were kept until the end of the experiment for each animal. The determination of serum creatinine was performed by the buffered kinetic reaction of Jaffe (diagnostic system of Boehringer Mannheim) with a Roche / Hitachi 917 system. Serum urea determination was performed by extracting 3 ml of blood with heparin to extract plasma (GD kinetic UV, Human, No. 10521). Measures were taken with a Cobas Mira Plus®6 autoanalyzer and a Hitachi 4020® spectrophotometer.

### Assessment of renal injury by histological examination

All animals underwent baseline renal biopsies followed by subsequent biopsies just after ischemia and one-week after injury. Samples were prepared by 10% formalin fixation and paraffin embedding, followed by staining with hematoxylin and eosin and Periodic acid–Schiff. A blinded pathologist using standard light microscopy assessed the degree of lesions at the tubular and interstitial level of all biopsy samples. The epithelial tubular affectation was scored as follow: 0: absence or dilation with reduction of the brush border; 1: proximal vacuolization with some isolated necrotic cell; 2: proximal vacuolization with disseminated necrotic cells; 3: proximal vacuolization with groups of necrotic cells. The Interstitial affection was score thereby: 0: absence of inflammatory infiltrate or <10% of parenchyma; 1: inflammatory infiltrate 10-25% of the parenchyma; 2: Inflammatory infiltrate 25-50% of the parenchyma; 3: Inflammatory infiltrate> 50% of the parenchyma.

### Microarray experiment

RNA was extracted from the PR, PS and WL kidney biopsies from each animal. The extractions were performed starting from 50 mg of each biopsy performed with the NZyol Kit following manufacturer instructions (Nzytech genes & enzymes). Microarray hybridization was carried out at High Technology Unit (UAT) at VHIR. RNA integrity was assessed by Agilent 2100 Bioanalyzer (Agilent, Palo Alto, Ca). Only samples with similar RNA integrity number were accepted for microarray analysis. Gene Titan Affymetrix microarray platform and the Genechip Porcine Gene 2.1 ST 16-Array plate were used for this experiment. This array analyzes gene expression patterns on a whole-genome scale on a single array with probes covering many exons on the target genome, and thus permitting expression summarization at the exon level or gene level. Starting material was 200 ng of total RNA of each sample. Briefly, sense ssDNA suitable for labeling was generated from total RNA with the GeneChip WT Plus Reagent Kit from Affymetrix (Affymetrix, Santa Clara, CA) according to the manufacturer’s instructions. Sense ssDNA was fragmented, labeled and hybridized to the arrays with the GeneChip WT Terminal Labeling and Hybridization Kit from the same manufacturer.

### Microarray data analysis

All microarray data in this publication have been deposited in NCBI’s Gene Expression Omnibus (49, 50) and are accessible through GEO Series accession number GSEXXXX (http://www.ncbi.nlm.nih.gov/geo/query/acc.cgi?acc=GSEXXXXX). Bioinformatic analysis was performed at the Statistics and Bioinformatics Unit (UEB) at VHIR. Robust Multi-array Average (RMA) algorithm (51) was used for pre-processing microarray data. Background adjustment, normalization and summarization of raw core probe expression values were defined so that the exon level values were averaged to yield one expression value per gene. The analysis was done considering the experimental factors (time points and sex) and taking into account the pairing between samples in most of the comparisons performed. Data were subjected to non-specific filtering to remove low signal and low variability genes. Conservative thresholds were used to reduce possible false negative results. This yields a list of 3435 genes to be analyzed. Selection of differentially expressed genes was based on a linear model analysis with empirical Bayes modification for the variance estimates (52). This method is similar to using a ‘t-test’ with an improved estimate of the variance. To account for multiple testing, P-values were adjusted to obtain stronger control over the false discovery rate (FDR), as described by the Benjamini and Hochberg method (53). Genes with adjusted P-value below 0.05 and absolute value of log2 fold change over 1 were called differentially expressed.

### Quantitative Reverse-transcription polymerase chain reaction (qRT-PCR)

Microarray experiments were validated by qRT-PCR experiments. Up to 2 µg of total RNA was retro-transcribed using the High Capacity RNA-to-cDNA Master Mix (Applied Biosystems) and used to perform quantitative gene expression analyses using TaqMan® Gene Expression Master Mix (Applied Biosystems). qPCR was performed in a 7900HT Fast Real Time PCR system (Applied Biosystems, Inc.) These specific TaqMan probes were used: *IFIT3* (Ss04248506_s1); *FABP5* (Ss03392150_m1); *CXCL10* (Ss03391845_g1); *CD274* (Ss03391947_m1) and *RSAD2* (Ss03381589_u1). To confirm the use of equal amounts of RNA in each reaction, all samples were examined in parallel for beta-actin (Ss03376160_u1). Triplicate PCR amplifications were performed for each sample. mRNA levels of human ischemic kidney biopsies were also measured at the same conditions. Non-tumoral post-ischemic renal tissues from 10 men and 9 women (36 to 83 years old) undergoing nephrectomy for renal cancer treatment were collected after 30 minutes of ischemia, thus corresponding to the post-surgery (PS) condition in our pig model. Specific TaqMan probes were used: *IFIT3* (Hs01922752_s1); *FABP5* (Hs02339439_g1); *CXCL10* (Hs00171042_m1) *CD274* (Hs00204257_m1) and *RSAD2* (Hs00369813_m1) for qRT-PCR experiments. To confirm the use of equal amounts of RNA in each reaction, all samples were examined in parallel for beta-actin (Hs01060665_g1).

### Ingenuity Pathway Analysis (IPA)

Ingenuity Pathway Analysis (IPA) to study the microarray data was conducted using the Qiagen software (https://digitalinsights.qiagen.com/). In our study, IPA was used to detect and overlap the most significant regulated genes across different time and sex comparisons. A distinct data-filtering criterion was set: a log fold change cut-off of ± 0.5. An individual analysis was performed for each comparison and the top 10 genes up- and down-regulated were reported. Their expression was represented in heatmaps.

### Gene Set Enrichment Analysis (GSEA) based pathway enrichment analysis

Pathway enrichment analysis was carried out by searching for enriched gene sets (e.g. pathways, molecular functional categories, complexes) for the different microarray comparisons using GSEA as previously described (54) and depicted in Figure Supp3. The pathway gene set definition (GMT) files loaded on the software were created with the archived instance of g:Profiler (Ensembl 93, Ensembl Genomes 40 (rev 1760, build date 2018-10-02) with a p-value cutoff: 0.01 for the microarray files. We used “gene set permutation” with 200 permutations to compute p-values for enriched gene-sets, followed by GSEA’s standard multiple testing correction.

### Enrichment Map pathway analysis visualization

The resulting enrichment results were visualized with the Enrichment Map plugin for the Cytoscape network visualization and analysis software. We loaded GSEA individual dataset using a FDR threshold between 0.01 and 0.1. A multi-dataset enrichment map comprising all comparisons was created with a FDR of 0.25. In the enrichment maps, each gene set is symbolized by a node in the network. Node size corresponds to the number of genes comprising the gene-set. The enrichment scores for the gene-set are represented by the node’s color (red indicates up-regulation, blue represents down-regulation). To identify redundancies between gene sets, the nodes are connected with edges if their contents overlap by more than 50%. The thickness of the edge corresponds to the size of the overlap. The 3.7.1 Cytoscape version (55) was used with the following apps: EnrichmentMap (54, 56), clusterMaker2 (57), WordCloud, (58), NetworkAnalyzer(59) and AutoAnnotate (60). Pathways are shown as circles (nodes) that are connected with lines (edges) if the pathways share many genes. Nodes are colored by ES, blue and red meaning down and up-regulated pathways, respectively. Edges are sized on the basis of the number of genes shared by the connected pathways. Network layout and clustering algorithms automatically group similar pathways into major biological themes. The EnrichmentMap software takes as input a text file containing pathway enrichment analysis results and another text file containing the pathway gene sets used in the original enrichment analysis. Gene sets were also visualized by heatmaps using online Heatmapper tool (61).

### Statistics

Results were expressed as the mean ± standard error of the mean (SEM). Student’s t-test (two-tailed) was used for statistical analysis. A P value of less than 0.05 was considered to indicate statistically-significant differences. Statistical analyses were made with commercially available software (GraphPad Prism, version 6.00 for Windows, GraphPad Software, La Jolla California USA). Bioinformatic Analysis was performed using the free R and Bioinformatic software.

## Author Contributions

SN, LC, AM, and JM contributed to conception and experimental design; LC performed IRI surgeries; SN, LC, DR, MES, MA, AS MF, JM and AM performed data acquisition or data analysis; and SN, GCR and AM prepared the manuscript, incorporating comments from other authors.

## Acknowledgements

We thank all members of the Renal Physiopathology Group for valuable discussions. DNA microarray and the bioinformatics analysis were carried out by the Statistics and Bioinformatics Unit (UEB) of the Vall d’Hebron Research Institute (VHIR). This work reflects only the authors’ views, and the EU Community is not liable for any use that may be made of the information contained therein.

## FIGURE LEGENDS

**Supplementary figure 1**. Number of genes differentially expressed throughout renal IRI in porcine kidney males and females at different time points (adjusted p-value ≤0.25 were considered. (PR: pre-ischemia; PS: post-ischemia; WL: one week later).

**Supplementary figure 2**. Number of genes differentially expressed throughout renal IRI in porcine kidney between males and females at the same time point (adjusted p-value ≤0.25 were considered. (PR: pre-ischemia; PS: post-ischemia; WL: one week later).

**Supplementary figure 3. Protocol overview of GSEA analysis**. Gene lists derived from diverse omics data undergo pathway enrichment analysis, using GSEA, to identify pathways that are enriched in the experiment. Pathway enrichment analysis results are visualized and interpreted in Cytoscape using its EnrichmentMap, AutoAnnotate, WordCloud and clusterMaker2 applications.

## TABLES LEGENDS

**Table 1**. Summary of clusters and gene sets included in the pattern 1 for the renal IRI time comparison.

**Table 2**. Summary of clusters and gene sets included in the pattern 2 for the renal IRI time comparison.

**Table 3**. Summary of clusters and gene sets included in the pattern 3 for the renal IRI time comparison.

**Table 4**. Summary of clusters and gene sets included in the pattern 4 for the renal IRI time comparison.

**Table 5**. Summary of clusters and gene sets included in the pattern 5 for the renal IRI time comparison.

**Table 6**. Summary of clusters and gene sets included in the pattern A for the renal IRI sex comparison.

**Table 7**. Summary of clusters and gene sets included in the pattern B for the renal IRI sex comparison.

**Table 8**. Summary of clusters and gene sets included in the pattern C for the renal IRI sex comparison.

**Table 9**. Summary of clusters and gene sets included in the pattern D for the renal IRI sex comparison.

**Table S1**. Serum creatinine and BUN levels in males and females pigs throughout the experiment

**Table S2**. Top 10 up- and down-regulated gene sets and NES in GSEA analysis for FPS vs FPR comparison

**Table S3**. Top 10 up- and down-regulated gene sets and NES in GSEA analysis for FWL vs FPS comparison

**Table S4**. Top 10 up- and down-regulated gene sets and NES in GSEA analysis for FWL vs FPR comparison

**Table S5**. Top 10 up- and down-regulated gene sets and NES in GSEA analysis for MPS vs MPR comparison

**Table.S6**. Top 10 up- and down-regulated gene sets and NES in GSEA analysis for MWL vs MPS comparison

**Table S7**. Top 10 up- and down-regulated gene sets and NES in GSEA analysis for MWL vs MPR comparison

**Table S8**. Top 10 up- and down-regulated gene sets and NES in GSEA analysis for MPR vs FPR comparison

**Table S9**. Top 10 up- and down-regulated gene sets and NES in GSEA analysis for MPS vs FPS comparison

**Table S10**. Top 10 up- and down-regulated gene sets and NES in GSEA analysis for MWL vs FWL comparison

